# Telomerase inhibition by MST-312 sensitises breast cancer cells to anti-cancer properties of Plumbagin

**DOI:** 10.1101/2023.05.29.542688

**Authors:** Safoura Sameni, Ramya Viswanathan, Gavin Yong-NG Quan, Wilner Martinez-Lopezm, Prakash Hande

## Abstract

Breast cancer is the most common cause of malignancy and the second most common cause of cancer death in women. This heterogeneous disease is currently broadly classified as ER, PG positive luminal tumours, HER2 amplified tumours and triple-negative breast cancers (TNBC). Natural plant derived compounds are proven to be promising anti-cancer chemotherapeutics agents with minimal cytotoxic effects on healthy cells. Plumbagin (5-hydroxy-2-methyl-1, 4-naphthoquinone) is a phytochemical derived from the roots of *Plumbago zeylanica* and it is known to possess anti-cancer properties similar to other compounds of naphthoquinones. In about 90 % of cancer cells, the telomerase enzyme activity is revived to add telomeric repeats to evade apoptosis. In this study, a combinatorial approach of combining anti-cancer compound Plumbagin to induce genotoxicity and a potent telomerase inhibitor, MST-312 (synthetic derivative of tea-catechins) was used to determine the synthetic lethality in breast cancer cells such as MDA-MB-231 (TNBC) and MCF-7 (lumina) cells. MDA-MB-231 cells were responsive to combination treatment to both short-term (48 hours) and long-term treatment (14 days) in a synergistic manner, whereas in MCF-7, the combination treatment was more effective in the long-term regimen. Furthermore, the cytotoxic effects of the Plumbagin and MST-312 combination treatment were not recoverable after the short-term treatment. In conclusion, combination treatment of MST-312 and Plumbagin is proven to be more effective than single Plumbagin compound treatment, in inducing DNA damage and telomere dysfunction leading to greater genome instability, cell cycle arrest and eventually cell death in cancer cells.

## Introduction

Telomerase is broadly expressed in most of human cancers, yet undetectable in somatic cells, suggesting an exciting target. Telomerase inhibition has been shown to boost the response to anti-cancer therapeutics in human breast cancer cells (Shay and Wright 2002, Shay and Wright 2005, Cerone, Londono-Vallejo et al. 2006, Venkatesan, Khaw et al. 2017, Sugarman, Zhang et al. 2019). In our previous study, telomerase inhibitory effects of Plumbagin were demonstrated upon long term chronic treatment of 32 days (Sameni and Hande 2016). We hypothesise that Plumbagin may exert a more effective targeted therapy with a shorter timespan in combination with a telomerase inhibitor through its implementation of its DNA damaging potential in breast cancer cells. We used MST-312, a synthetic derivative of tea-catechin, which was reported to inhibit the telomerase enzyme activity (Seimiya, Oh-hara et al. 2002). MST-312 blocks the cell cycle at the G2/M phase, increases pH2AX protein level, and induces apoptosis in lung cancer cells. MST-312 also provokes G2/M cell cycle arrest and acute ATM-pathway-dependent DNA damage in breast and brain cancer cells (Wong, Ma et al. 2009, Gurung, Lim et al. 2014, Morais, Arcanjo et al. 2019). Therefore, we sought to investigate the anti-cancer effects of combinatorial approach on targeting telomerase by MST-312 and Plumbagin treatment, in TNBC MDA-MB-231 and luminal MCF-7 breast cancer cells as compared to the relative normal MCF-10A cell type. These combinatorial experiments were set for short-term (48 hours) and long-term (14 days) studies to inspect the cell viability, DNA damage, telomere/telomerase equilibrium, cell morphology, clonogenicity, cell cycle, immunofluorescence, genome stability and protein expression levels of these cells, upon dual treatments.

## Materials and Methods

### Cells

Two human mammary epithelial carcinoma cell types, MDA-MB-231, known as a typical triple negative breast cancer cell type and MCF-7, as an ER+/PG+ luminal tumour, were obtained from American Type Culture Collection (ATCC, Rockville, MD. USA). These were grown in RPMI-1640 (Roswell Park Memorial Institute medium) (Gibco), supplemented with 10% foetal bovine serum (FBS) and 0.5% penicillin/streptomycin. These two breast cancer cell types have formed the breast cancer models of this study. All cells were grown and incubated at 37°C in a humidified 5% CO_2_ incubator. All the supplements used were from Gibco, USA, unless otherwise stated.

### Drug Treatment

Plumbagin (Sigma-Aldrich, Missouri, USA) was dissolved in 100 % sterile dimethyl sulfoxide (DMSO) (Sigma, USA) to prepare 100 mM stock solution from which, working concentrations were prepared later. One µl of the working concentrations in 0.0, 1.0, 2.0, 3.0 and 4.0 mM were added in the cell culture to achieve 0.0, 1.0, 2.0, 3.0 and 4.0 μM final concentrations, respectively. Short term studies were performed following 48 hours treatment with different concentrations of Plumbagin, mainly at the IC_50_ dose. As for chronic long-term studies, they were carried out for either 14 or 32 days with 2/3^rd^ of IC_50_ doses. This range of concentration has been reported to be practicable *in vivo* and with low or no toxicity to normal cells (Hsieh, Lin et al. 2006, Padhye, Dandawate et al. 2012, Khaw, Sameni et al. 2015, Sameni and Hande 2016).

Telomerase inhibitor, MST-312 (Merck, USA), was dissolved in sterile DMSO to prepare the stock solution of 10 mM and thereafter it was more diluted to achieve different working concentrations. Appropriate amounts of these stocks were added to the cultured cells to obtain the desired final concentrations. Twenty-four hours prior to exposure of cells to Plumbagin, cells were treated with 0.5 and 1 μM of MST-312 for long-term and short-term treatments, unless otherwise stated. Both mentioned concentrations are known to be non-toxic to normal cells (Seimiya, Oh-hara et al. 2002, Gurung, Lim et al. 2014).

### Crystal Violet Assay

This assay was performed to verify how MST-312 and/or Plumbagin affect the density of cells by modulating their cell growth or inducing cell death. Growing cells were treated with a range of drug concentrations, following which the inhibitory concentration at 50% cell viability (IC_50_) was obtained and selected for further studies. The details of this assay were described in our earlier (Khaw, Sameni et al. 2015, Sameni and Hande 2016). Cells were treated with MST-312, Plumbagin and/or in combination of both drugs for 24 hours at different concentrations as indicated and cell viability was measured in treated cells.

### Cell Morphology and Colony Formation Assay

For morphological observations, light microscopy was utilized. After treatment of cells with appropriate agents, images of cell morphology were captured at 40X, 100X and 200X magnifications of microscope, using an Olympus C-7070 WZ digital camera (Japan).

To determine the effects of drug treatments on the colony formation ability, clonogenic or colony formation assay was performed. In four independent sets, cells were seeded and after 48 hours treatment, 2 x 10^3^ cells were allowed to grow for 14 days in fresh medium. Colonies were then stained with crystal violet solution (Sigma, USA) (as described above), and dried at room temperature. Thereafter large colonies (> 50,000 cells) were detected and counted as survived number of colonies.

### Cell cycle analysis by flow cytometry

Cell cycle analysis was performed by fluorescence-activated cell sorting (FACS) methodology. Following treatment with drugs, cells were collected, fixed in 70% ethanol: 1X PBS, resuspended in 0.4 mL of 1xPBS containing propidium iodide (PI) (Sigma, USA), RNase A (Roche, USA) and 0.1% Triton X (Biorad, USA); (2 mg propidium iodide and 2 mg RNaseA/100 mL 1X PBS), incubated in the dark for 30 minutes at 37°C. For each treatment, two independent sets were used and for each 10,000 events per sample were analysed using flow cytometry (FACSCalibur™, Becton Dickinson, USA) at 488 nm excitation λ and 610 nm emission λ. Obtained data was analysed using Summit 4.3 software (Beckman Coulter, USA).

### Immunofluorescence staining for γH2AX

To specifically identify the amount of double strand breaks (DSB) triggered by Plumbagin treatment, γH2AX protein as an indicator of this phenomenon was assessed. Cells were seeded on 12mm cover slips in six well plates, treated with Plumbagin, washed in PBS and fixed in 4% paraformaldehyde for 15 minutes in an orbital shaker. Following permeabilization with 0.2% of Triton-X-100 for 10 minutes at 4°C, cells were blocked in 5% BSA: PBS and immunostained with anti-γH2AX (Ser 139) (Millipore., USA) for 1 hour. After washing with PBS, FITC-conjugated anti-mouse antibody (eBioscience, USA) was applied to each slide. Then the incubation for 1 hour in the dark was succeeded by washing three times with 1X TBS-T, 15 minutes each. The nuclei were counterstained by DAPI solution with mounting media (Vectashield®, Vector Laboratories, Burlingame, CA, USA). Fluorescent images of 150 randomly selected samples of three different experimental set were obtained by confocal microscopy (Olympus Fluoview FV1000).

### Peptide nucleic acid-fluorescence in-situ hybridisation (PNA-FISH)

Cells were arrested at mitosis and chromosome preparations were made as described previously. Slides with metaphase spreads were subjected to PNA-FISH to determine telomere specific chromosome changes following treatments with MST-12 and Plumbagin, either in single treatment or in combination. The details of PNA-FISH and chromosome analysis were explained in our earlier publications (Poonepalli, Balakrishnan et al. 2005, Gurung, Lim et al. 2014, Sameni and Hande 2016, Zeegers, Venkatesan et al. 2017) Microscopic analysis was performed using Zeiss Axioplan 2 Imaging fluoresnce microscope and images were analysed using in situ imaging software (Metasystems, Germany).

### Immunofluorescence staining for γH2AX and telomere dysfunction induced foci (TIF)

Cells were seeded on 12 mm cover slips in 6-well plates and treated for 48 hours. After the treatments, cells were washed in PBS and fixed in 4% paraformaldehyde for 15 minutes in an orbital shaker. Following permeabilization with 0.2% of Triton-X-100 for 10 minutes at 4°C, cells were blocked in 5% BSA: PBS and immunostained with anti-γH2AX (Ser 139) (Millipore, USA) for 1 hour. After washing with PBS, FITC-conjugated anti-mouse antibody (eBioscience, USA) was added to the slides for 1 hour. Thereafter, the washing was conducted with TBS-T, 4 % formaldehyde for 15 minutes and 0.2 % Triton-X-100, samples were hybridised with Cy3-labelled telomere sequence-specific PNA probes by first denaturing at 80°C for 6 minutes and incubation for 2 hours in dark at room temperature. Following hybridisation, the slides were washed in formamide and 10 % Tween-20 for 5 minutes. Finally, the cells were stained with DAPI (Vectashield®). Approximately one hundred images were captured on fluorescence microscope (Carl Zeiss Axioplan ǁ, Germany) with triple filters for simultaneous observation of DAPI (blue), FITC (green) and Cy3 (red).

### Protein expression and western blot studies

After drug exposure, the total cellular proteins of control and treated cells were isolated using lysis buffer (distilled water containing 50 mM Tris HCL, pH 8, 5mM EDTA, 0.15 M sodium chloride, 0.5% NP40, 0.5 mM DTT, 1× PhosSTOP (Phosphatase inhibitor cocktail tablets, Roche, Germany), 1X Protease cocktail tablet (Roche, Germany), following which lysates were rotated at 4°C at 14,000 rpm for 30 minutes. Protein concentration was determined by the colorimetric Bradford assay (Bio-Rad,USA) with bovine serum albumin (BSA) as a standard. Western blot analyses for proteins involved in different pathways, including cell proliferation, cell cycle regulatory proteins (p21, p53, Cyclin B1, Cyclin D), apoptosis related agents (caspases 8 and 9), DNA damage response pathways (Phospho-H2AX (Millipore, USA), PARP, ATM (GeneTex, USA), Phospho-ATM (GeneTex), DNA-PKcs, Phospho-DNA-PKcs and telomere-telomerase equilibrium (hTERT (Epitomics, USA), POT1, TRF1, TRF2).Antibodies were purchased from Santa Cruz Biotechnology, USA, unless otherwise stated. For different protein expression profiles, the experiments were repeated between two to six times.

### Telomerase activity detection using Telomerase Repeat Amplification Protocol (TRAP)

To measure the telomerase activity, TRAP assay was carried out using TRAPeze® XL Telomerase Detection Kit (Millipore, USA). All steps were done according to the manufacturer’s instructions. Fluorescence signals of PCR products were measured using fluorescence plate reader (TECAN SpectraFluor Plus, Männedorf, Switzerland).

### Telomere length measurement by Terminal Restriction Fragment (TRF)

Following treatment with drugs, the total DNA was extracted by DNeasy Tissue Kit (Qiagen, USA). To measure the telomere length, the TeloTAGGG Telomere Length Assay kit (Roche Molecular Systems, USA) was utilized. The chemiluminescence telomere signals were developed onto x-ray films and the telomere lengths were calculated and analysed by the Molecular Imaging Systems software (Kodak MI application, Eastman©, V4.00, USA).

### Statistical analysis

Statistical significance of the difference between and among the groups was assessed by Student’s t-test using Microsoft Excel (Microsoft Corporation, USA) and two-way ANOVA using GraphPad Prism (version 6.00 for Windows, GraphPad software, San Diego California USA, www.graphpad.com). P values lower than 0.05 were considered statistically significant (P ˂0.05). Wherever applicable, the mean values are expressed as mean ± standard deviation (SD) from two or three independent experiments.

## Results

### MST-312 treatment slightly reduces cell proliferation capacity

The crystal violet cell viability assay was performed to assess the toxicity of MST-312 to cells after 24 hours. Figure 1A shows a concentration dependant drop in cell intensity of breast cancer and normal cells, signifying dose dependant toxicity. Nevertheless, in breast cancer cells, the overall cell survival rate was not significantly affected in the presence of MST-312 for the range of 0-5 µM after 24 hours. Initially, to ascertain the appropriate dose and time point for telomerase inhibition (Gurung et al. 2014), a dose response study (0 to 5 µM) was performed for 24-hour treatment with MST-312 in MCF10A, MCF-7 and MDA-MB-231 cells (Figure 1A). Breast cancer cells displayed higher sensitivity to MST-312 compared to MCF10A cells.

**Figure 1.**
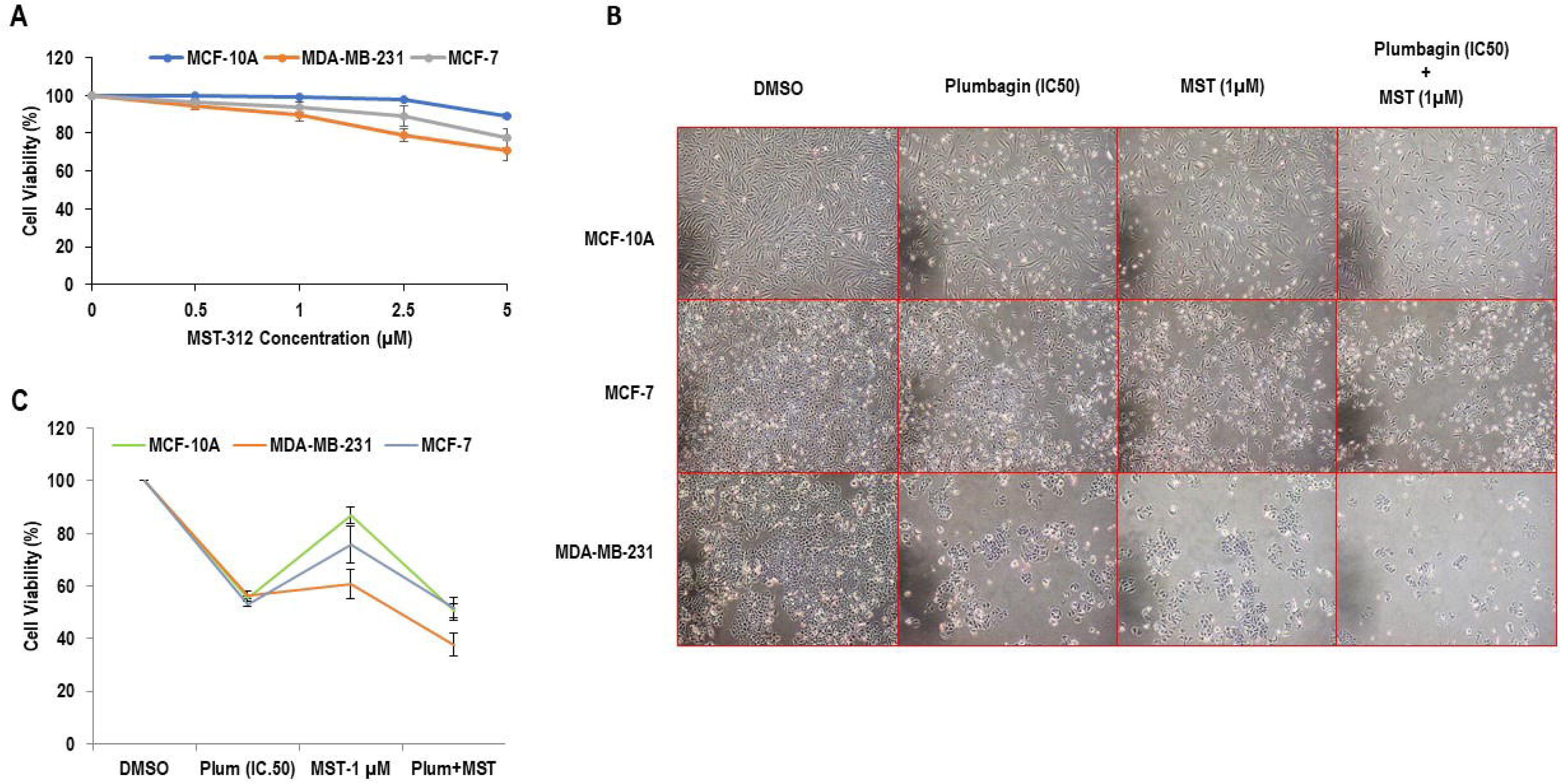
**A** Cytotoxicity of MST-312 in breast cancer cells and MCF-10A. After 24 hours of treatment with MST-312, cells were subjected to crystal violet cell viability assay. The Y-Axis represents the percentage of reduction in cell survival compared to control. **B – C**: Telomerase inhibition limits cell proliferation in Plumbagin treated cancer cells. Cells were pre-treated with MST-312 (1 μM) for 24 hours followed by Plumbagin (IC_50_) treatment for another 48 hours. Thereafter, they were subjected to: **(B**) cell morphology by light microscopy with the magnification of 100X showing cell death and lower cell density after dual treatment, and (**C**) crystal violet assay on the same cells indicated more reduction in cell survival upon co-treatment of Plumbagin and MST-312 in MDA-MB-231 cells. The percentage of cell viability was standardized against DMSO control. “Plum” denotes Plumbagin, “MST” signifies MST-312.

According to the data from this study and our previous study (Gurung, Lim et al. 2014, Gurung, Lim et al. 2014) which demonstrated the relatively lower toxicity but efficient inhibition of telomerase (Gurung, Lim et al. 2014) up to 1 µM dose of MST-312, this dose was considered as the main dose of MST-312 in most of the subsequent studies.

### Telomerase inhibition sensitises the breast cancer cells to Plumbagin-induced cell death

To examine the effect of Plumbagin, combined with the telomerase inhibitor MST-312 on the survival of breast cancer cells, morphological observation and crystal violet assays were conducted (Figure 1 B and C). Cells were pre-incubated with MST-312 (1 µM) for 24 hours prior to Plumbagin (IC_50_ doses) exposure to ensure that telomerase inhibitor has exerted its effects before Plumbagin becomes effective. The percentage of cell viability was normalised to results from non-exposed cells (control). With Plumbagin treatment alone, the viability of cells reduced to about 53 % for respective IC_50_ doses of each cell type, validating the IC_50_ doses selected in previous section. MST-312 treatment of 1 µM reduced the viability of the cells by 13 % in MCF10-A, 40 % in MDA-MB-231 and 25 % in MCF-7. In combination treatment, there was about 50% reduction in MCF-7 and MCF-10A, and 67 % reduction in MDA-MB-231 cells, with more floating and rounded features, as the manifestations of dying cells. The cytotoxic effect of dual treatment of MCF-7 and MCF-10A cells was similar to that of Plumbagin single treatment. Interestingly, reduction in MDA-MB-231 cell viability was 20% greater in dual treatment; compared to Plumbagin single treatment.

To explore the clonogenic ability of telomerase inhibited cells after Plumbagin treatment, colony formation essay was conducted in MDA-MB-231 and MCF-7 breast cancer cells. After 24 hours of pre-treatment with MST (1 µM) and a subsequent 48-hour treatment with Plumbagin (IC_50_), the cells were allowed to recuperate in drug-free medium for 14 days. Only the cells which survived the treatment were able to form colonies. Decrease in cell proliferation in colonies could be due to either cell cycle arrest or apoptosis caused by the treatment. Plumbagin reduced the colony formation ability in both cancer cell types by 41 %, and 18% in MDA-MB-231 and MCF-7, respectively when compared to DMSO (Figure 2A and B). However, MST-312 treatment decreased the number of colonies by 70 % and 72 % in MDA-MB-231 and MCF-7 cells, respectively, in comparison to DMSO controls. A decrease of 72 % in MDA-MB-231 and 86% in MCF-7 cells colonies over the combination treatment, suggested that MST-312 treatment impaired the colony formation ability of cells upon Plumbagin treatment.

**Figure 2:**
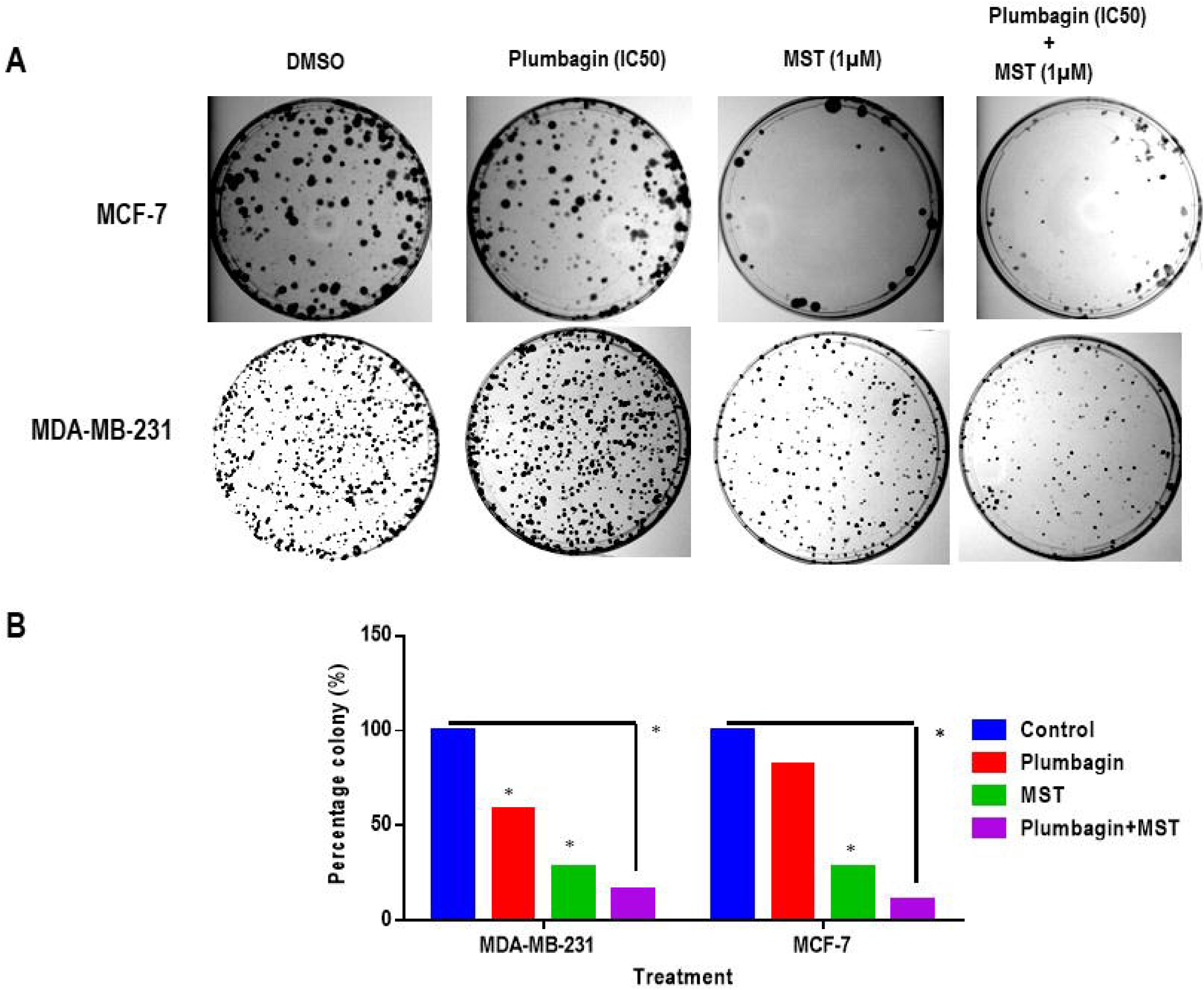
Plumbagin-MST-312 co-treatment decreased the clonogenic ability of cells. **(A)** In the colony formation assay, only drug-resistant cells are able to proliferate and form colonies after treatment. The rest of the cells are either arrested or underwent apoptosis progressively within the 14 days recovery period following the 48 hours treatment with Plumbagin (IC_50_) and MST-312 (1μM). **(B)** Representative image of colony formation capacity of breast cancer cells treated with Plumbagin and MST-312. (*) indicates P <0.05 when compared to DMSO control.

### Telomerase inhibition produces significant synergistic apoptosis in breast cancer cells, when combined with Plumbagin

Microscopic observation of cells treated with both telomerase inhibitor (MST-312) and Plumbagin showed indication of massive cell death as compared to mono-treatments. Presence of large number of floating cells after combined incubation period together with lower clonogenicity and survival rate in breast cancer cells implies a potential for cell apoptosis. Hence, in order to evaluate the cell growth situation of combined treatments, cell cycle studies were conducted using FACS methodology. Cell cycle assay was done after 14 days of treatment with Plumbagin (2/3^rd^ of IC_50_) and MST-312 (MDA-MB-231: 0.5 µM, MCF-7: 1 µM). The results confirmed breast cancer cell apoptosis through the observed increase in the sub-G1 population (Figure 3A). These observations also led to the other inference that MST-312 promotes both cells towards G2/M arrest. Interestingly, in MDA-MB-231, both Plumbagin and MST-312 exerted apoptotic effects, showing that combination treatment is synergistic. Notably, mono-treatment with MST-312 in MCF-7 cells potentiated G2/M arrest more than Plumbagin treatment. However, combinational treatment of both MST-312 and Plumbagin resulted in an upregulation of both sub-G1 and G2/M levels.

**Figure 3:**
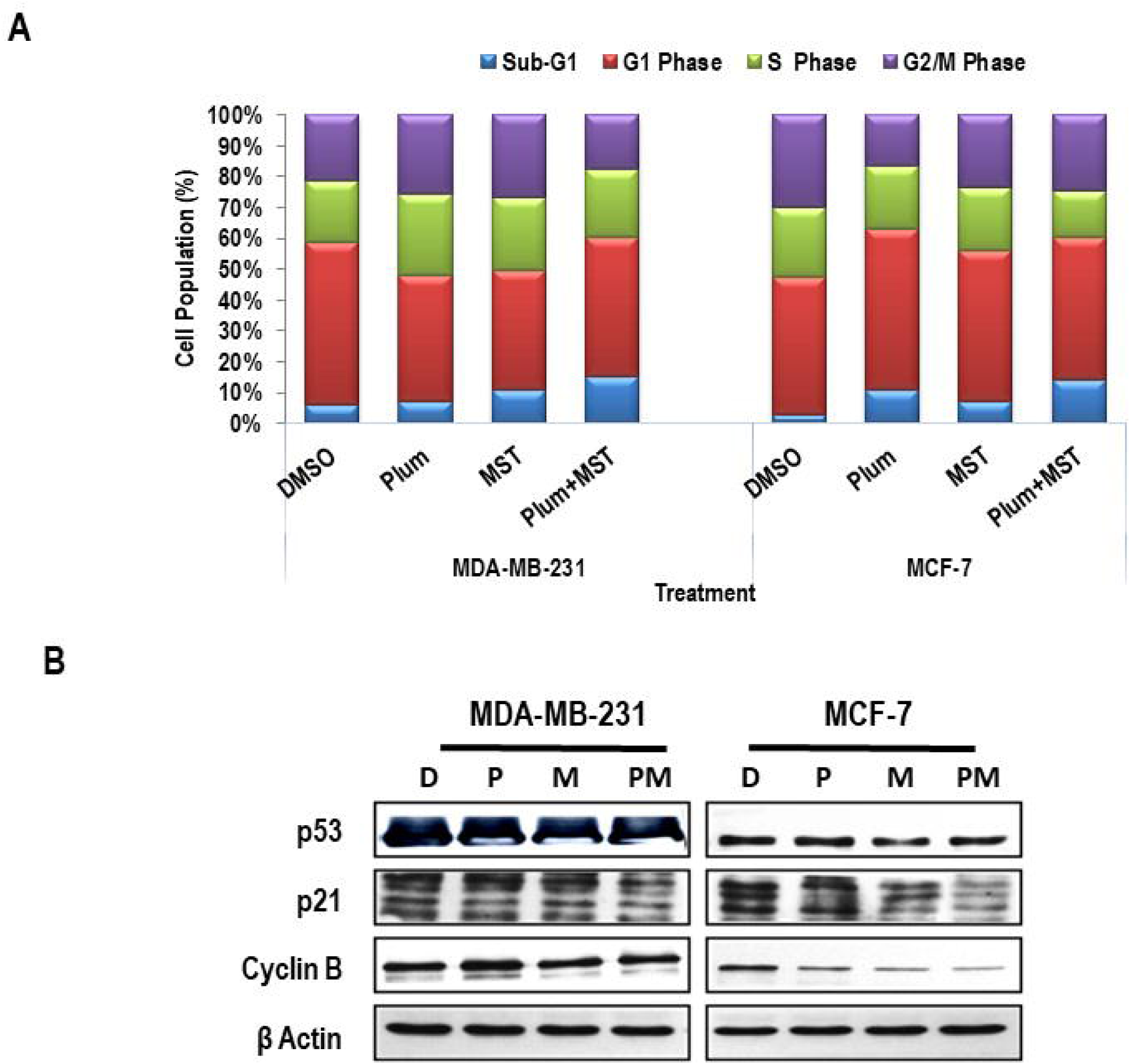
Co-treatment of Plumbagin and MST-312 induces growth arrest and apoptosis in breast cancer cells. (A) Cell cycle profile of 14 days treatment with Plumbagin (2/3^rd^ of IC_50_) and MST-312 (MDA-MB-231: 0.5μM and MCF-7: 1μM) in breast cancer cells, using propidium iodide staining and flow cytometry detection. Though MST-312 potentiates cell cycle arrest at G2/M phase, population of apoptotic cells is greater in combined treatment than in individual treatments. (*) indicates P <0.05 when compared to DMSO. (B) Changes in level of cell cycle regulatory proteins were also evaluated. Whole cells were lysed, and equal amounts of proteins were separated using 10-12% SDS-PAGE, transferred to PVDF membrane and immunoreacted with antibodies against p53, p21 and Cyclin B. β-actin was used as loading control. “Plum” denotes Plumbagin, “MST” signifies MST-312. D: DMSO control; P: Plumbagin; M: MST-312; PM: Plumbagin+MST-312

To validate cell cycle data, immunoblotting of respected proteins was performed (Figure 3B). The protein expression of Cyclin B1, as the indicative of G2/M arrest was reduced in both cells, and substantially in MCF-7, after dual treatment with MST-312 and Plumbagin. To determine the possible involvement of p53 in inducing growth arrest, the expression of p53 and p21 was evaluated after all types of treatments. The level of p21 was reduced in MCF-7 cells after MST-312 and subsequently more after combination treatment, showing greater sensitivity towards MST-312 and Plumbagin combined treatment.

Taking these findings together, cell cycle analysis demonstrated that MST-312 treatment arrested the cells mainly at G2/M as compared to Plumbagin, and the combined treatment of these two drugs led to increased synergistic cell death.

### MST-312 reduces the telomerase activity and shortens telomere length in Plumbagin-treated breast cancer cells, without impairing the hTERT protein expression

To evaluate the effectiveness of telomerase inhibitor MST-312 on telomerase activity, TRAP assay was performed. The percentage of telomerase activity was shown to be interrupted by both mono-treatment and combined treatments of Plumbagin and MST-312 (Figure 4A). In MDA-MB-231 cells, there was a reduction in telomerase activity by 21 %, 24 %, and 44 % in comparison to DMSO control, correspondingly in Plumbagin, MST-312 and combination treatments. It can be interpreted that reduction in telomerase activity in combination is contributed to both telomerase inhibitor MST-312 and Plumbagin. In MCF-7 cells, telomerase activity decreased by 26 % (Plumbagin), 77 % (MST-312), and 84 % (combination) compared to DMSO control, which shows that reduction in telomerase activity in combination treatment is mostly contributed to telomerase inhibitor MST-312.

**Figure 4:**
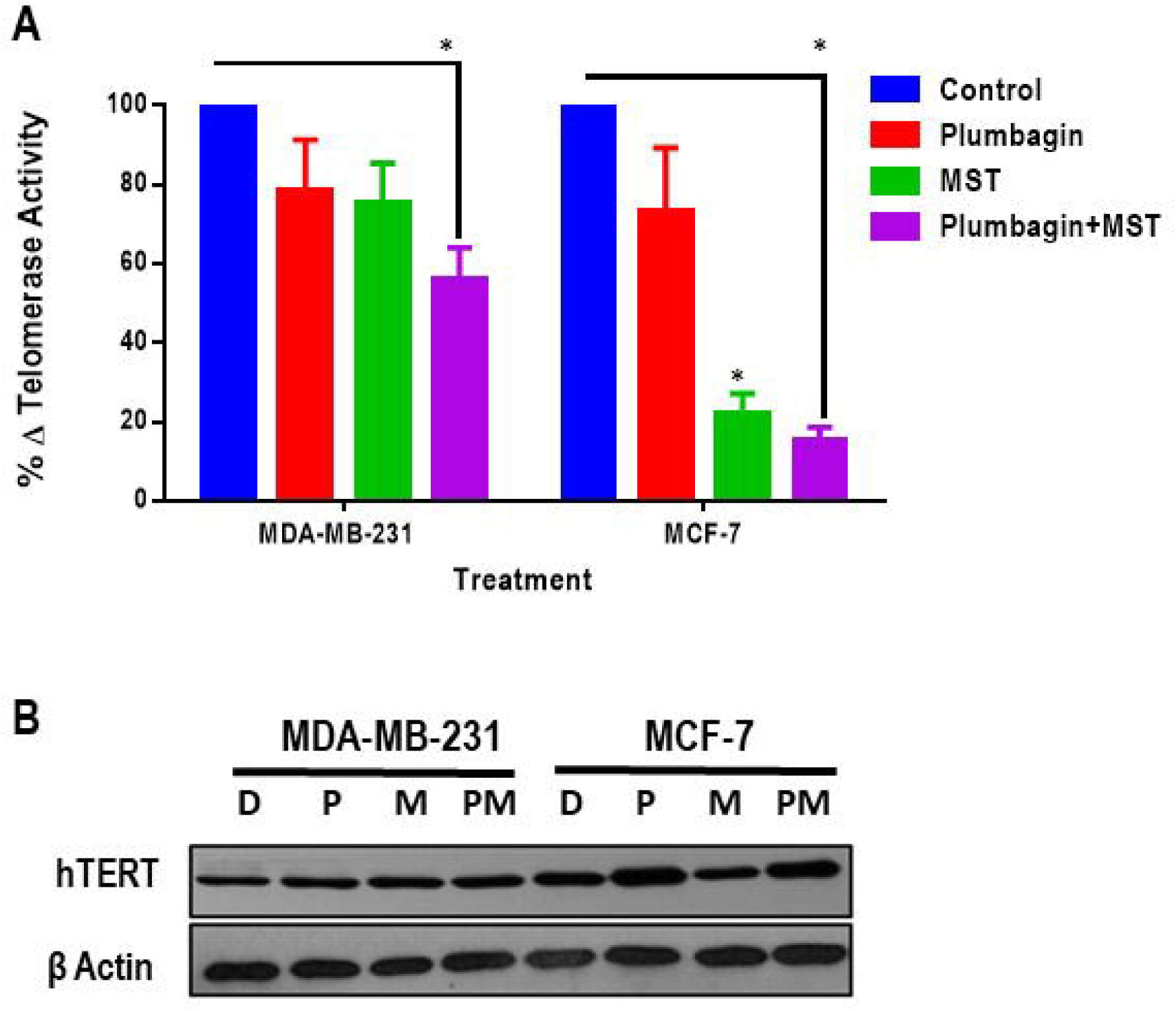
A. MST-312 efficiently reduces the telomerase activity in breast cancer cells. Following 14 days of treatment with Plumbagin (2/3^rd^ of IC_50_) and MST-312 (0.5 μM for MDA-MB-231 and 1 μM for MCF-7), the telomerase activity was detected by TRAP (telomeric repeat amplification protocol) assay. In MCF-7 combination treatment, more reduction in the percentage of telomerase activity was detected. (*) indicates P <0.05 when compared to DMSO control. B. **MST-312 does not decrease the expression level of hTERT, the catalytic subunit of telomerase.** hTERT expression level was determined compared to β-actin as the loading control, after 14 days of treatment with Plumbagin (2/3rd of IC_50_) and MST-312 (MDA-MB-231-0.5μM and MCF-7-1μM). D: DMSO control; P: Plumbagin; M: MST-312; PM: Plumbagin+MST-312.

Decrease in telomerase activity can be due to interruption of the function of telomerase enzyme and/or down-regulation of the expression of telomerase enzyme. Even though there was a decrease in telomerase activity as seen in TRAP results, hTERT expression (which is the catalytic subunit of telomerase enzyme) was not down-regulated (Figure 4B). hTERT expression level in MDA-MB-231 and MCF-7 cells had no significant change in Plumbagin and combination treatment. This result indicates that telomerase inhibition by MST-312 did not involve the downregulation of hTERT expression level.

To investigate if the inhibition of telomerase activity was translated into reduction of telomere length, terminal restriction fragment analysis was done. The cells were treated for 14 days with Plumbagin (2/3^rd^ of IC_50_) and MST-312 (MDA-MB-213: 0.5 µM and MCF-7: 1μM). Telomere length reduction was observed across both breast cancer cell types following telomerase inhibition (Figure 5A and B). In MDA-MB-231 cells, the telomere length of control cells, treated with DMSO, was observed as 2444 bp and this length was further reduced by 112 bp in Plumbagin treated cells and 438 bp in co-treated cells after 14 days. In MCF-7 telomere length decreased by 309 bp in Plumbagin treated cells and 1164 bp in combination treatment compared to DMSO treated controls. Reduction in telomere length was 3-fold greater in MCF-7 combination treatment in comparison to control cells.

**Figure 5:**
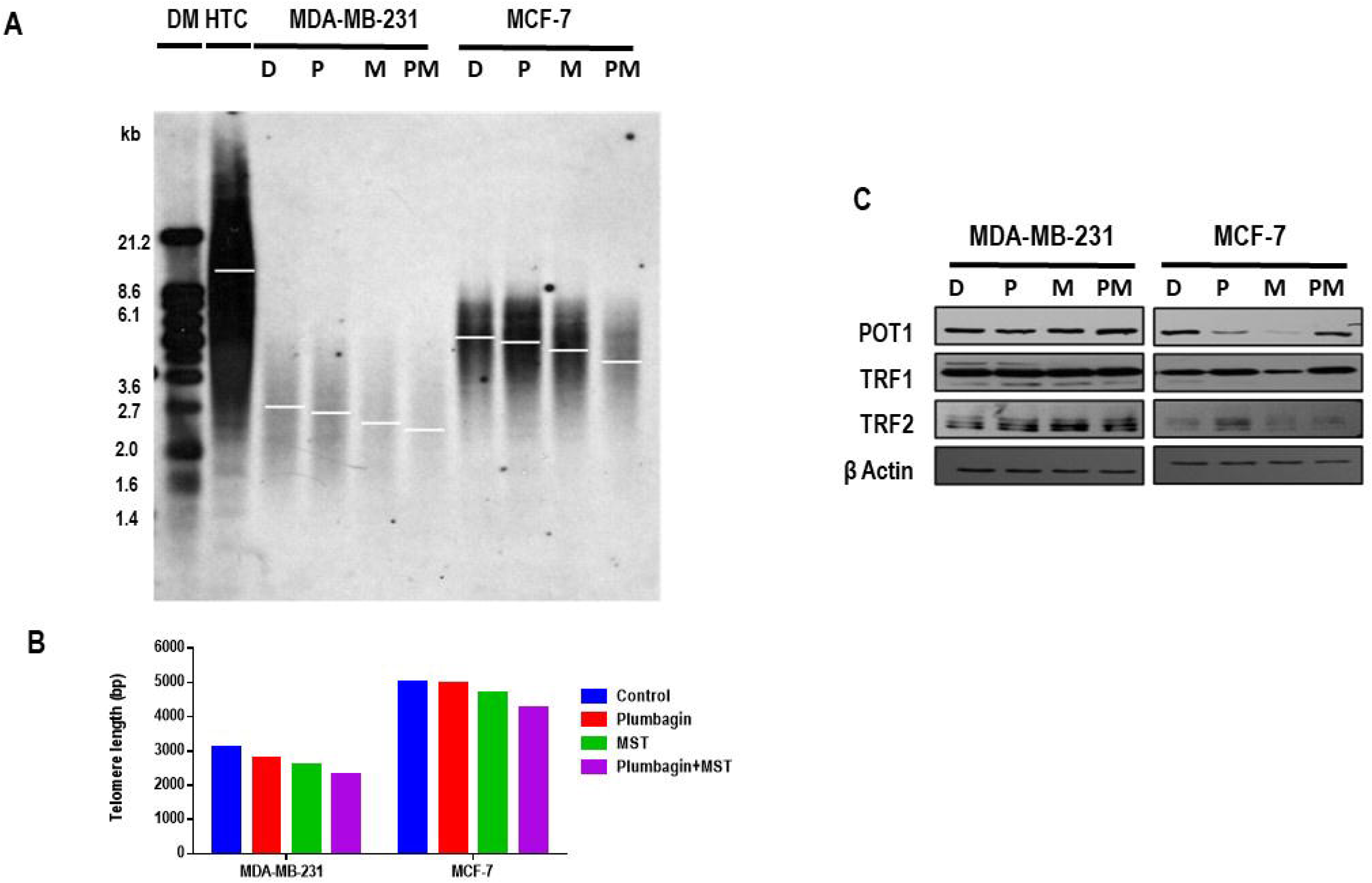
Co-treatment of Plumbagin and MST-312 induces more telomere shortening in cancer cells. Terminal Restriction Fragment (TRF) measuring telomere length after 14 days of treatment with Plumbagin (2/3^rd^ of IC_50_) and MST-312 (MDA-MB-213: 0.5 μM and MCF-7: 1μM). (A) Southern blot analysis of telomeric regions. Analysis was done using Kodak imaging software. (B) Graphical presentation of the telomere length of samples compared to DIG marker (DM) and the high telomere control (HTC). (C) Western blot analysis on TRF1, TRF2 and POT1 proteins upon treatment. β-actin was used as loading control. D: DMSO control; P: Plumbagin; M: MST-312; PM: Plumbagin+MST-312

Telomere-associated proteins such as TRF1, TRF2 and POT1 are important in telomere-telomerase dynamics, the expression levels of these proteins were investigated in the same treated cells (van Steensel, Smogorzewska et al. 1998, Smogorzewska, van Steensel et al. 2000, Baumann and Price 2010, Maciejowski and de Lange 2017, Sugarman, Zhang et al. 2019). Reduction in telomere length of MST-312-treated cells was corroborated by decreased protein expression levels of POT1, TRF1 and TRF2 through western blot in MCF-7 cells (Figure 5C). On the other hand, for MDA-MB-231 cells, there is no significant changes for POT1 and TRF1, but a slight up-regulation in the expression level of TRF2 was observed in the order of mono- and co-treatments, as compared to the control cells (Figure 5C). The reduction in telomere length could most probably be due to telomerase inhibition by MST-312 in combination treatments rather than Plumbagin’s effects after 14 days.

Taken together, these data highlighted that dual treatment with telomerase inhibition and Plumbagin induces the telomere attrition in MCF-7 and MDA-MB-231 cells, more than the Plumbagin treatment alone.

### Inhibition of telomerase in breast cancer cells results in elevated Plumbagin-induced DNA damage and telomere dysfunction

To investigate the extent of DSB caused by Plumbagin (IC_50_) and MST-312 (1 µM) treatment across DNA, as well as its specificity to telomere region after 48 hours of treatment, telomere dysfunction induced foci (TIF) analysis was performed (Figure 6A and B). Presence of DSB was further validated through western blot analysis of γH2AX, with β-actin as reference. In TIF, DSB was represented by green γH2AX foci, telomere regions were denoted by red telomere probe and co-localisation (TIF) of both signals resulted in yellow dots.

**Figure 6:**
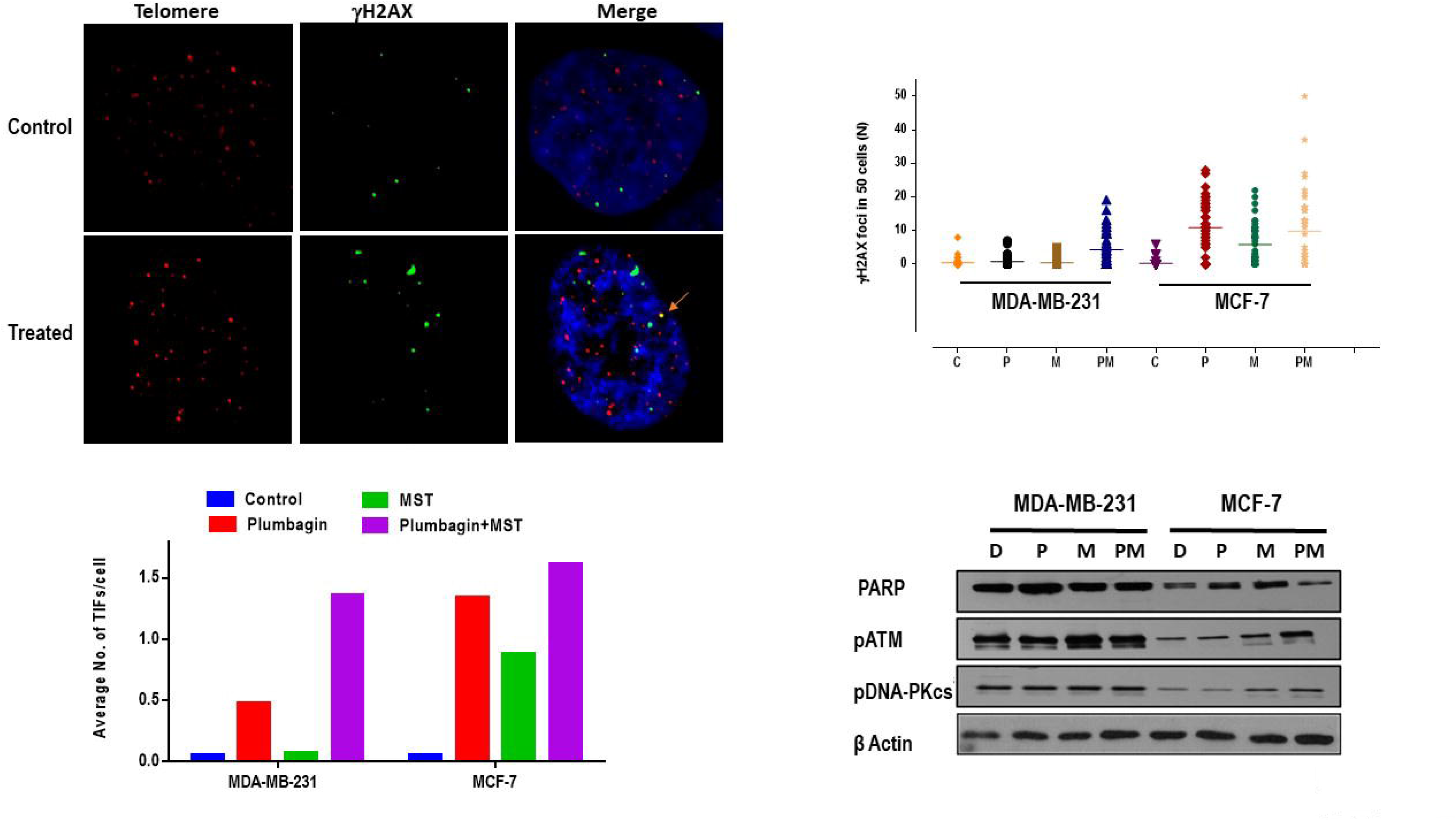
Induction of telomere dysfunction following telomerase inhibition in Plumbagin treated cells. Colocalization of ɤH2AX and telomeres represents telomere dysfunction induced foci (TIF) after being treated for 48 hours with Plumbagin (IC_50_) and MST-312 (1μM). (A) Immunofluorescence image of TIF in control and treated cells. Bright yellow dots are visible in treated cells with colocalization. (B) Average number of TIFs per cells.

To detect the frequency of DSB occurring at telomeric regions, number of co-localized foci was counted. Average number of TIFs per cell in MDA-MB-231 upon Plumbagin treatment was 0.5-fold greater than untreated DMSO control cells, whereas in MCF-7, it was 1.3 times greater than DMSO (Figure 6B). Upon MST-312 treatment the number of TIFs in MDA-MB-231 cells was not significant, though it increased by 0.8-fold in MCF-7 cells. In combination treatment, number of TIFs was 1.4 and 1.8-fold higher than in control cells in MDA-MB-231 and MCF-7, respectively; inferring that telomerase inhibition enhances the probability of DSB incidences at the telomeres.

### MST-312 promotes the extent of Plumbagin-induced double strand breaks in human breast cancer cells

Cells were treated with Plumbagin (IC_50_) for 48 hours, followed by 24-hours pre-incubation with MST-312 (1μM). Thereafter immunoblotting and immunocytochemistry detection of γH2AX was employed to measure the protein expression and average number of γH2AX foci, as a representative of double strand breaks. **(C)** Average number of γH2AX foci per cell, indicating presence of double strand breaks across the genome. **(D)** Western blot analysis of γH2AX protein expression level as compared to the β-Actin standard control. D: DMSO control; P: Plumbagin; M: MST-312; PM: Plumbagin+MST-312. **E. MST-312 modulates the DNA damage response pathway proteins in breast cancer Plumbagin-treated cells.** Representative blots for PARP, pATM and pDNA-PKcs proteins following 24-hours pre-incubation with MST-312 (1μM) and 48 hours treatment with Plumbagin (IC_50_). β-actin was used as loading control. Protein level alterations upon dual treatments were more evident in MCF-7 cells, compared to MDA-MB-231 cells. D: DMSO control; P: Plumbagin; M: MST-312; PM: Plumbagin+MST-312

Similar trend was observed for treatment-induced DBS (Figure 6B), indicating the average number of γH2AX foci in Plumbagin and combination treatment for both cell types. In MDA-MB-231 cells, there was an insignificant increase in the average DNA damage upon MST-312, while there were about 5-fold increases in the average number of γH2AX foci for MCF7 cells. With respect to Plumbagin treatment, in MDA-MB-231 cells, the increase of γH2AX foci in combination treatment was 4-fold higher while interestingly, in MCF-7 cells this number declined by one. The findings of this study indicated that the DNA damage observed in combination treatment mostly contributed by Plumbagin in short term treatment.

To validate the observations in TIF assay, western blot analysis was performed for γH2AX protein expression level. As compared to the corresponding β-Actin, the trend of γH2AX protein expression levels soar from Plumbagin to the combination treatment in both cancer cell types (Figure 6D. This was in line with the trends observed in TIF assay as seen previously. In consistent with the TIF findings and the trend of γH2AX foci, the phosphorylation level of γH2AX protein was less upon MST-312 and combined treatments in MCF-10A cells, suggestive of the fact that MST-312 might attenuate the DNA damaging potential of Plumbagin in normal cells (data not shown).

Once again, the observations of TIF together with the telomere attrition and telomerase inhibition were more altered in response to MST-312 and accordingly in MST-312 and Plumbagin combined treatments in both cells highlighting that the mechanism of action of Plumbagin is more oriented towards exerting DNA damage to cancer cells, while MST-312 affects the telomere dynamics mechanisms.

To further examine the effects of telomerase inhibition on DNA damaging abilities of Plumbagin in breast cancer cells, immunoblotting was carried out for other key proteins involved in DDR pathways. The enhanced DNA damaging potential of Plumbagin in combined treatments was corroborated by higher expression levels of phosphorylated DNA-PKcs and phosphorylated ATM (Figure 6D). In MCF-7 cells, PARP-1 expression level was increased in either of MST-312 and Plumbagin treatments, but when combined the protein expression seemed to be attenuated. Whereas expression level of pDNA-PKcs increased in MST-312 and combination treatments compared to single exposure to Plumbagin. On the other hand, the expression levels of PARP-1 and pDNA-PKcs were relatively constant in MDA-MB-231 control, Plumbagin, MST-312, and combination treatments. For the pATM protein expressions, there was a slight up-regulation in the expression levels of this protein for both breast cancer cells, after 48 hours of single and co-treatment with MST-312, when compared to Plumbagin treatment (Figure 6D).

Together the findings of this study confirm the earlier reports establishing the fact that MST-312 acts through activation of the ATM/pH2AX DNA damage pathway in short-term treatments of cancer cells (Serrano, Bleau et al. 2011).

### Telomerase inhibition of human breast cancer cells promotes the amount of Plumbagin induced chromosomal instability

As demonstrated in our earlier study (Sameni and Hande 2016), Plumbagin exposure resulted in telomere dysfunction after 32 days. It is speculated that the inhibition of telomerase in breast cancer cells may lead to enhanced Plumbagin-induced chromosomal instability within 14 days; therefore PNA-FISH was performed to investigate genomic instability and chromosomal aberrations. Following 14 days of chemical inhibition of telomerase by MST-312 (MDA-MB-231: 0.5 µM and MCF-7: 1 µM) and Plumbagin treatment (2/3^rd^ of IC_50_) in breast cancer cells, telomere associated aberrations were detected such as telomere signal loss, chromosome fusion without telomere signals (dicentric and tricentric chromosomes), multiple telomere signals, and also chromosome or chromatid breaks. There were about 12 %, 16 %, 17 % and 22 % total chromosomal aberrations observed in control, Plumbagin, MST-312, and combination treatments, respectively in MCF-7 cells. Whereas, in MDA-MB-231 cells, correspondingly the total aberrations were 3 %, 5 %, 6 % and 8 % in control, Plumbagin, MST-312, and combination treatments. In both the cell types, the percentages of chromosomal aberrations were larger in combined exposures than in Plumbagin (single treatment group) after 14 days of treatment (Figure 7A-C and Table 1).

**Figure 7.**
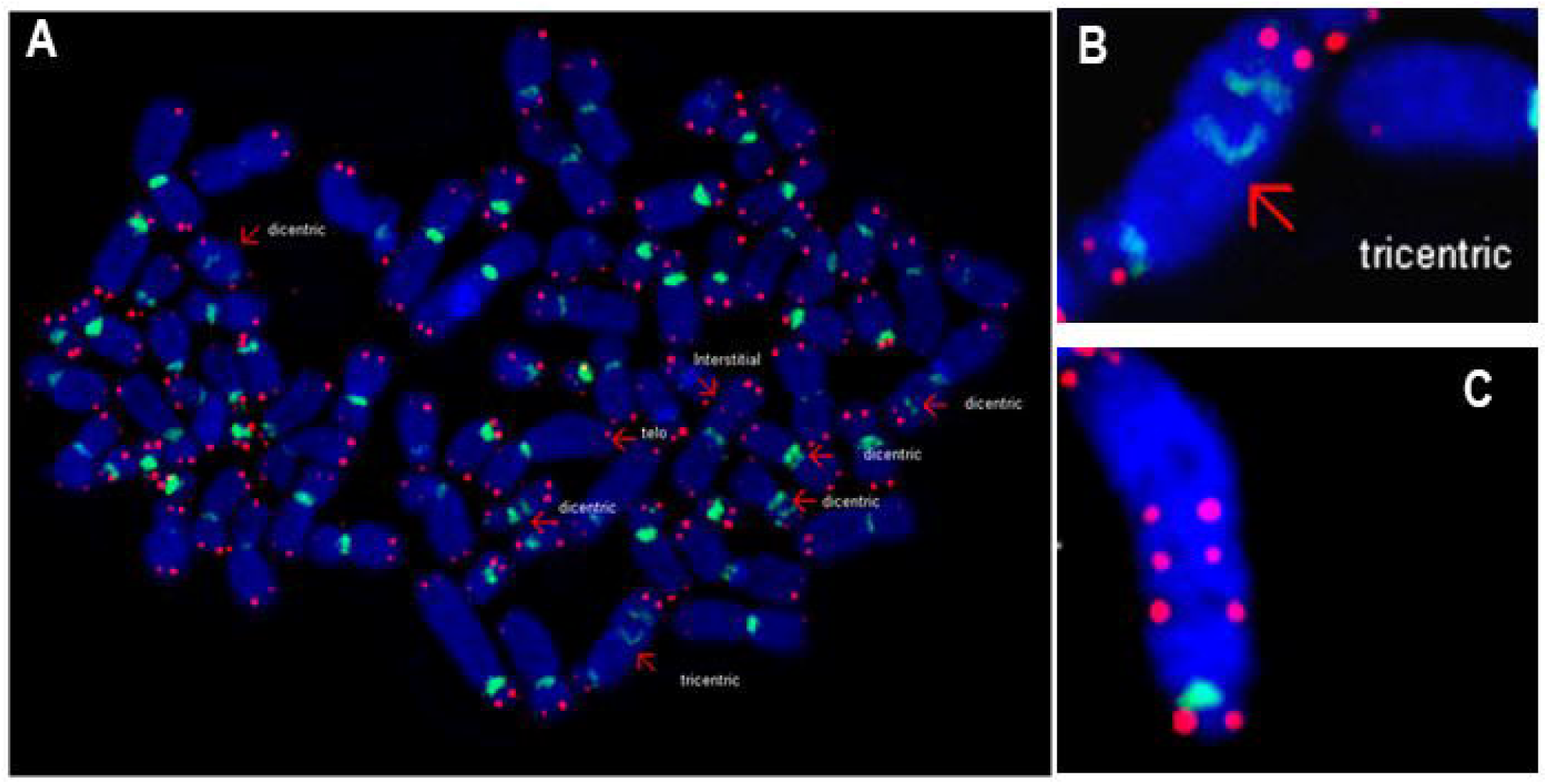
A – C: Effects of combined treatment with MST-312 (telomerase inhibition) and Plumbagin on the chromosome stability. Peptide Nucleic Acid-Fluorescence in-Situ Hybridisation (PNA-FISH) was performed on breast cancer cells treated with Plumbagin (2/3^rd^ of IC_50_) followed by pre-incubation with MST-312 (MDA-MB-231: 0.5μM and MCF-7: 1μM). The findings are evidence of increase in chromosomal aberrations of dual treatments after 14 days of treatment. Plum: Plumbagin; MST: MST-312.

**Table 1:** Chromosomal changes in breast cancer cells following treatment with plumbagin, MST and in combination.

| Cell lines | Treatments | No. of Metaphases | Average chromosome number ± SD | Ends without telomeres | End-to-end Fusions | Breaks | % Total aberrant cells |
| --- | --- | --- | --- | --- | --- | --- | --- |
| MCF-7 | DMSO | 20 | 61 ± 8.9 | 0 | 5 | 1 | 12% |
| | Plum | 20 | 59 ± 4.6 | 6 | 4 | 4 | 16% |
| | MST | 20 | 60.7 ± 5.4 | 7 | 4 | 6 | 17% |
| | Plum+MST | 20 | 58.5 ± 8.7 | 10 | 2 | 1 | 22% |
| MDA-231 | DMSO | 20 | 50.2 ± 5.9 | 3 | | | 3% |
| | Plum | 20 | 47.5 ± 6.9 | 3 | | 2 | 5% |
| | MST | 20 | 51 ± 2.5 | 2 | 1 | 3 | 6% |
| | Plum+MST | 20 | 41.8 ± 8.7 | 6 | 1 | 1 | 8% |

## Discussion

Anticancer activity of Plumbagin against human breast cancer cells, including inhibition of cell growth and apoptosis induction, was shown to be partly due to inhibition of NF-kB/Bcl-2 pathway (Padhye, Dandawate et al. 2012). Throughout the combination studies on MST-312 and Plumbagin treatment, interestingly, it was found that Plumbagin exerts efficient anti-cancer effects on the highly resistant and metastatic MDA-MB-231 breast cancer cells. In MDA-MB-231 cells, the combination treatment of Plumbagin with MST-312 had noteworthy effects in both short term and long-term studies. Treatment with MST-312, a telomerase inhibitor, sensitised the MDA-MB-231cells to Plumbagin treatment following both short term and long-term exposure. This could probably be due to the shorter basal telomere length in MDA-MB-231 cells compared to MCF-7 as observed in control results. It was reported by Hemann and colleagues that it is not the average telomere length, but rather the critically short telomeres, which trigger cellular responses to the loss of telomere function (Hemann, Strong et al. 2001). The lag phase, which is required to critically shorten the telomere length, would be less for shorter telomeres. Furthermore, based on the immunocytochemistry observations, Plumbagin treatment showed greater genotoxic consequences in MDA-MB-231 cells. Plumbagin possibly affects cell density by inhibiting cell growth or inducing cell death as observed in cell cycle assay. In long term treatment, Plumbagin and MST-312 single treatments caused G2/M phase arrest while Plumbagin and MST-312 combination treatment more obviously led to increased apoptosis.

In MCF-7 cells, the combination treatment with MST-312 yielded almost equal cytotoxic effects as Plumbagin single treatment in short term. This was observed in both cell viability and TIF experiments, where the observed cytotoxic and genotoxic effects in the combination treatment could also be contributed to the presence of Plumbagin. In these short-term studies, the less sensitization of MCF-7 cells towards MST-312 treatment could be due to longer lag phase which MCF-7 requires because of its longer telomere length. Noteworthy, after 14 days of treatment, MCF-7 cells were more affected in combination treatment than single treatments with Plumbagin. This indicates that MST-312 has begun to show its effects along with Plumbagin in combination treatments after 14 days. It was reported by Seimiya and colleagues (Seimiya, Oh-hara et al. 2002) that when cells are treated with telomerase inhibitor, those with longer telomeres would require more doubling time to exhaust telomeric sequence compared with cells with shorter telomeres. As reported earlier (Sameni and Hande 2016), MCF-7 cells have longer population doubling time. Furthermore, TRF findings also showed longer basal telomere length for MCF-7 cells, which collectively might point to one of the main reasons for the difference of cellular responses of MDA-MB-231 and MCF-7 cells to combination treatments in short term and long-term studies. Furthermore, both cell types are known to be genetically distinct in many ways (Sameni and Hande 2016) (data not shown). Therefore, the response to combination treatment varies between the cell types and it is time dependent. MCF-7 cells require longer time than MDA-MB-231 cells to respond in combination treatment compared to Plumbagin treatment alone. Though there is longer lag phase observed in MCF-7 cells due to longer telomere length, Plumbagin continue to induce DNA damage, making the combination treatment more efficient.

The cytotoxic effects of Plumbagin and MST-312 combination treatments appeared to be irreversible, unlike that of Plumbagin single treatments as discussed earlier. Only cells that were resistant to the Plumbagin and MST-312 treatments during short term exposure were able to form colonies. Other cells were either arrested or underwent apoptosis. However, the results indicated that very few cells were able to form colonies in combination treatment in both cell types, even though the treatment was terminated after 48 hours, and the cells were allowed to recuperate in drug-free medium for 14 days. This shows that the inflicted damage by combination treatments were too extensive that the cells were not able to repair or overcome the cell cycle arrest. Such deleterious damages could not only have targeted DNA but also caused damage to other cellular components (Roos and Kaina 2006, Brinkman, Liu et al. 2021).

Reduction in telomerase activity was mostly contributed to telomerase inhibition in both cells. In MCF-7, the reduction in telomerase activity has led to telomere attrition. Depletion in the telomerase activity rate of MCF-7 cells was five folds from the control cells to combined treated ones with Plumbagin and MST-312, though the rate of telomere shortening was slower. When telomerase activity is inhibited, cancer cells would face the end replication problem similar to normal cells due to absence of telomerase activity.

As manifested by the results, when cells were treated with Plumbagin and MST-312 single treatments, they were arrested at G1 phase in MCF-7 cells, which ensured that damaged DNA would not replicate. This infers why the number of cells undergoing cell division in Plumbagin and MST-312 co-treated cells were much lower than that of controls. This might also explain how the rate of telomere length attrition was not commensurate with rate of telomerase activity reduction in MCF-7 cells. In long term treatments of MCF-7 cells, the reduction in telomere length, though slow, but together with the relevant cytotoxic effects were significant enough as evident by the increase in TRF1 and POT1 protein expression levels in combination treatments. POT1 is a negative regulator of telomerase activity (Kelleher, Kurth et al. 2005, Gu, Jia et al. 2021). TRF1 negatively regulates telomere length and it was reported that overexpression in TRF1 (Munoz, Blanco et al. 2009) or overproduction of wild type TRF1 induces telomere shortening (van Steensel and de Lange 1997, Iwano, Tachibana et al. 2004, Maciejowski and de Lange 2017, Nassour, Schmidt et al. 2021). POT1 binds the 3’-overhang, whereas TRF2 directly binds the double-stranded telomeric DNA; and both protect telomeres from DDR by representing the ATM and ATR kinase signalling pathways, respectively.

In both MCF-7 and MDA-MB-231 cells, single treatment with Plumbagin showed up-regulation in the expression levels of TRF2. This may be due to DNA damaging properties of Plumbagin proving the information that upon DNA damage, telosome reacts by up-regulating the TRF2 expression and telomerase activity in malignant cells. It was shown that this up-regulation of TRF2 protein expression might act as antiapoptotic mechanisms in the DDR of malignant cells (Klapper, Qian et al. 2003). Telomerase inhibition by MST-312 single treatment did not modify the TRF2 expression level in MCF-7 cells [in this study and (Gurung, Lim et al. 2014)]; though in MDA-MB-231 cell type, it resulted in the increased trend of TRF2, both at the single exposure or as combined with Plumbagin treatment. This might be due to the pure epithelial nature of MDA-MB-231 breast cancer cell types (Cailleau, Young et al. 1974), since it was verified that telomerase abrogation accelerates TRF2-induced epithelial carcinogenesis (Blanco, Munoz et al. 2007). Furthermore, it has been illustrated that telomerase inhibition might not be effective to cease the growth of TRF2-overexpressing tumours (Blanco, Munoz et al. 2007). Both TRF1 and TRF2 were reported as negative regulators of telomere length and their down-regulation was shown to be important to maintain telomeric DNA in cancer (Yamada, Tsuji et al. 2002). Therefore, their up-regulation might disturb the telomere dynamics.

Regarding the genotoxicity and cytotoxicity of each drug, MCF-7 cells were mainly prone to Plumbagin, especially in short term studies, as it was evident in TIF and cell cycle sub-G1 population number, respectively. In these findings apoptosis was more observed in Plumbagin single treatments than MST-312 ones. However, synergistic effects of MST-312 and Plumbagin treatments after 14 days were noticeable; and might have also been more effective if the duration of studies was longer.

On the contrary, as for MDA-MB-231 cells, the genotoxicity and cytotoxicity of treatments were synergistically exerted by genotoxic effects of Plumbagin and telomere dysfunction through telomerase inhibition of MST-312. Telomerase involvement in long term maintenance of telomere length and chromatin structure might potentiate its response to genotoxic stimuli. It was reported by Tamakawa and colleagues that telomerase positive breast cancer cells, respond to genotoxic agents in a cell cycle stage–specific manner: telomerase activity can increase cell survival by alleviating the toxic effect of genotoxic agents that are predominantly active in G2 phase of the cell cycle (Tamakawa, Fleisig et al. 2010). Hence when DNA damage is induced in S/G2 phase of the cell cycle, telomerase inhibition increases the cytotoxicity of a genotoxic stimulus (Tamakawa, Fleisig et al. 2010). Furthermore, it has been reported earlier that telomerase inhibition increased the susceptibility of malignant glioblastoma cells to Cisplatin-induced apoptosis (Kondo, Kondo et al. 1998, Seimiya, Oh-hara et al. 2002). The findings of this study are consistent with previous reports (Kondo et al.1998 and Seimiya et al 2002 (Gurung, Lim et al. 2014)), were Plumbagin and telomerase inhibitor MST-312 treatments impacted greatly and synergistically on the telomerase activity inhibition, apoptosis rate and cell viability of MDA-MB-231 cells. Cell cycle assay showed that Plumbagin-induced DNA damage occurs at G2 phase. Accordingly, combining the telomerase inhibitory effects of MST-312 with Plumbagin increased the cytotoxicity and genotoxicity of the MDA-MB-231 cells in both short- and long-term studies.

PARP-1 response was up-regulated in Plumbagin-treated MCF-7 cells, but inert in MDA-MB-231 cells. However, the cellular reaction of PARP-1 towards MST-312 treatments was ruled out in both the cell types, as evidenced by the steady level of protein expression throughout the exposure. However, it was presented that the protein levels of both pATM and pH2AX were elevated upon combined treatment of Plumbagin and MST-312 in MDA-MB-231 and MCF-7 cells. The higher levels of these two proteins could be believed to be due to MST-312 impact to large extents, as it was evident in immunoblotting findings of this telomerase inhibitor single treatment. These observations substantiate the report that MST-312 performs through activation of the ATM/pH2AX DNA damage pathway in short term-treated cancer cells (Serrano, Bleau et al. 2011).

The total number of aberrations which was mainly due to telomere signal loss was corroborated by TIF assay. This association advocated that upon combination treatment with Plumbagin and MST-312, the DSB DNA damages were colocalized at the telomere sites leading to more critical cancer cell genome instability. Genotoxic effects of Plumbagin treatment led to DSBs across genome and specifically at telomeric regions. Moreover, with MST-312 combination, it was observed more telomere instability such as telomere signal loss, chromosomes with multiple telomere signals, and chromosome fusions without telomere signals resulting in dicentric and tricentric chromosomes. Telomere attrition followed by telomerase inhibition and DSBs at telomeric regions resulted in deprotection of chromosome ends and eventually chromosomal fusion. This could most likely be due to the fact that the dysfunctional telomere repeats were too short for the efficient binding of telomere associated proteins such as TRF2 (Broccoli, Smogorzewska et al. 1997). Short telomeres would then evoke DNA repair activities that would result in chromosome fusions, most likely to be mediated by NHEJ repair mechanism. Increase in phosphorylated DNA-PKcs expression level in MCF-7 cells and constant expression level in MDA-MB-231 cells was also observed. Phospho-DNA-PKcs participates in NHEJ repair mechanism as mentioned previously. Hence, more telomere related chromosome dysfunctions in MCF-7 cells were observed than in MDA-MB-231 cells.

The presence of dicentric or tricentric chromosomes, produced by telomeric fusions, poses problem to the cell for subsequent cell division (Londono-Vallejo 2008). Multiple centromeres might be brought to opposite poles during anaphase and the resulting bridges would be broken, leaving DSB at one end. The broken ends would then be fused to another chromosome with uncapped telomere, which would reinitiate a cycle of breakage-fuse-bridge (BFB) (Gisselsson, Pettersson et al. 2000, Maciejowski and de Lange 2017, Chakravarti, LaBella et al. 2021). As there are many chromosomes with dysfunctional telomeres, different chromosomes fuse together randomly. Repetition of BFB in pre-crisis state would lead to widespread genome disintegration with fusion of different chromosome regions, resulting in chromosome ploidy changes. Further replication would lead to greater number of unstable chromosomes; thus, the cells can no longer divide without losing vital genetic material. Cells then enter mitotic catastrophe and die (Macera-Bloch, Houghton et al. 2002). The two main sources of telomere fusion and telomere dysfunctions observed in PNA-FISH results were due to telomere erosion by MST-312 treatment, and DSB caused by Plumbagin specifically at telomere region as seen in TIF results. According to TIF, the average number of TIF per cell was much higher in combination treatment in both MDA-MB-231 and MCF-7 cells, which might have promoted the genome instability of breast cancer cells.

Decrease in telomerase activity can be due to interruption of the function of telomerase enzyme and/or down-regulation of the expression of telomerase enzyme. Though the telomerase activity was down-regulated by telomerase inhibitor MST-312, the expression level of hTERT continues to increase in both cell types. As described earlier, the ribonucleoprotein telomerase consists of telomerase reverse transcriptase subunit (TERT), as the catalytic subunit, and telomerase RNA (hTER) that provides template for synthesis of telomere. These two components need to be assembled in order to form the fully functional telomerase. However, the molecular mechanism by which hTERT and hTR are assembled into a functional ribonucleoprotein is unknown (Bachand, Kukolj et al. 2000, Roake and Artandi 2020). As the mechanism of action of MST-312 is becoming clearer (Seimiya, Oh-hara et al. 2002, Gurung, Lim et al. 2014, Gurung, Lim et al. 2014, Fernandes, Gala et al. 2023), we speculate that decrease in telomerase activity by MST-312 could be due to interruption of the function of telomerase enzyme or the assembly of the telomerase. MST-312 was not shown to cause telomere damage, rather inhibited the telomerase activity as observed in short term TIF.

Telomerase is present in the majority of cancer cells and not in normal cells, except for stem cells and germ lines. Hence, a dual treatment comprising telomerase inhibitor would more specifically target towards breast cancer cells to elicit genome instability and affect the telomere dynamics of the breast tumour cells. The combination treatment of two natural products, Plumbagin and MST-312, was observed to be highly effective in long term and short-term studies in TNBC cells MDA-MB-231. Whereas the cytotoxic effects of combination treatment were more apparent in MCF-7 cells following long term treatment. The required longer period and its efficacy upon Plumbagin single treatment in MCF-7 cells was due to the difference in lag phases between these two cell types, contributed to the difference in initial telomere length.

Moreover, in MDA-MB-231, Plumbagin and MST-312 combination yielded synergistic effects in terms of DNA damage and impairment at the telomeres and telomere dynamics. The cytotoxic effects of dual treatment with Plumbagin and MST-312 are not recoverable, unlike that of Plumbagin single treatment. In conclusion, the combination treatment of MST-312 and Plumbagin was more effective than Plumbagin single treatment in terms of DNA damage and telomere dysfunction, which might lead to greater genome instability and cell cycle arrest, and eventually cell death in cancer cells.

## References

1. Bachand, F., G. Kukolj and C. Autexier (2000). “Expression of hTERT and hTR in cis reconstitutes and active human telomerase ribonucleoprotein.” RNA 6(5): 778–784.

2. Baumann, P. and C. Price (2010). “Pot1 and telomere maintenance.” FEBS Lett 584(17): 3779–3784.

3. Blanco, R., P. Munoz, J. M. Flores, P. Klatt and M. A. Blasco (2007). “Telomerase abrogation dramatically accelerates TRF2-induced epithelial carcinogenesis.” Genes Dev 21(2): 206–220.

4. Brinkman, J. A., Y. Liu and S. J. Kron (2021). “Small-molecule drug repurposing to target DNA damage repair and response pathways.” Semin Cancer Biol 68: 230–241.

5. Broccoli, D., A. Smogorzewska, L. Chong and T. de Lange (1997). “Human telomeres contain two distinct Myb-related proteins, TRF1 and TRF2.” Nat Genet 17(2): 231–235.

6. Cailleau, R., R. Young, M. Olive and W. J. Reeves, Jr. (1974). “Breast tumor cell lines from pleural effusions.” J Natl Cancer Inst 53(3): 661–674.

7. Cerone, M. A., J. A. Londono-Vallejo and C. Autexier (2006). “Telomerase inhibition enhances the response to anticancer drug treatment in human breast cancer cells.” Mol Cancer Ther 5(7): 1669–1675.

8. Chakravarti, D., K. A. LaBella and R. A. DePinho (2021). “Telomeres: history, health, and hallmarks of aging.” Cell 184(2): 306–322.

9. Fernandes, S. G., K. Gala and E. Khattar (2023). “Telomerase inhibitor MST-312 and quercetin synergistically inhibit cancer cell proliferation by promoting DNA damage.” Transl Oncol 27: 101569.

10. Gisselsson, D., L. Pettersson, M. Hoglund, M. Heidenblad, L. Gorunova, J. Wiegant, F. Mertens, P. Dal Cin, F. Mitelman and N. Mandahl (2000). “Chromosomal breakage-fusion-bridge events cause genetic intratumor heterogeneity.” Proc Natl Acad Sci U S A 97(10): 5357–5362.

11. Gu, P., S. Jia, T. Takasugi, V. M. Tesmer, J. Nandakumar, Y. Chen and S. Chang (2021). “Distinct functions of POT1 proteins contribute to the regulation of telomerase recruitment to telomeres.” Nat Commun 12(1): 5514.

12. Gurung, R. L., H. K. Lim, S. Venkatesan, P. S. Lee and M. P. Hande (2014). “Targeting DNA-PKcs and telomerase in brain tumour cells.” Mol Cancer 13: 232.

13. Gurung, R. L., S. N. Lim, G. K. Low and M. P. Hande (2014). “MST-312 Alters Telomere Dynamics, Gene Expression Profiles and Growth in Human Breast Cancer Cells.” J Nutrigenet Nutrigenomics 7(4-6): 283–298.

14. Hemann, M. T., M. A. Strong, L. Y. Hao and C. W. Greider (2001). “The shortest telomere, not average telomere length, is critical for cell viability and chromosome stability.” Cell 107(1): 67–77.

15. Hsieh, Y. J., L. C. Lin and T. H. Tsai (2006). “Measurement and pharmacokinetic study of plumbagin in a conscious freely moving rat using liquid chromatography/tandem mass spectrometry.” J Chromatogr B Analyt Technol Biomed Life Sci 844(1): 1–5.

16. Iwano, T., M. Tachibana, M. Reth and Y. Shinkai (2004). “Importance of TRF1 for functional telomere structure.” J Biol Chem 279(2): 1442–1448.

17. Kelleher, C., I. Kurth and J. Lingner (2005). “Human protection of telomeres 1 (POT1) is a negative regulator of telomerase activity in vitro.” Mol Cell Biol 25(2): 808–818.

18. Khaw, A., S. Sameni, S. Venkatesan, G. Kalthur and M. Hande (2015). “Plumbagin alters telomere dynamics, induces DNA damage and cell death in human brain tumour cells.” Mutat Res Genet Toxicol Environ Mutagen 793: 86–95.

19. Klapper, W., W. Qian, C. Schulte and R. Parwaresch (2003). “DNA damage transiently increases TRF2 mRNA expression and telomerase activity.” Leukemia 17(10): 2007–2015.

20. Kondo, Y., S. Kondo, Y. Tanaka, T. Haqqi, B. P. Barna and J. K. Cowell (1998). “Inhibition of telomerase increases the susceptibility of human malignant glioblastoma cells to cisplatin-induced apoptosis.” Oncogene 16(17): 2243–2248.

21. Londono-Vallejo, J. A. (2008). “Telomere instability and cancer.” Biochimie 90(1): 73–82.

22. Macera-Bloch, L., J. Houghton, M. Lenahan, K. K. Jha and H. L. Ozer (2002). “Termination of lifespan of SV40-transformed human fibroblasts in crisis is due to apoptosis.” J Cell Physiol 190(3): 332–344.

23. Maciejowski, J. and T. de Lange (2017). “Telomeres in cancer: tumour suppression and genome instability.” Nat Rev Mol Cell Biol 18(3): 175–186.

24. Morais, K. D. S., D. D. S. Arcanjo, G. P. de Faria Lopes, G. G. da Silva, T. H. A. da Mota, T. R. Gabriel, D. D. A. Rabello Ramos, F. P. Silva and D. M. de Oliveira (2019). “Long-term in vitro treatment with telomerase inhibitor MST-312 induces resistance by selecting long telomeres cells.” Cell Biochem Funct 37(4): 273–280.

25. Munoz, P., R. Blanco, G. de Carcer, S. Schoeftner, R. Benetti, J. M. Flores, M. Malumbres and M. A. Blasco (2009). “TRF1 controls telomere length and mitotic fidelity in epithelial homeostasis.” Mol Cell Biol 29(6): 1608–1625.

26. Nassour, J., T. T. Schmidt and J. Karlseder (2021). “Telomeres and Cancer: Resolving the Paradox.” Annu Rev Cancer Biol 5(1): 59–77.

27. Padhye, S., P. Dandawate, M. Yusufi, A. Ahmad and F. H. Sarkar (2012). “Perspectives on medicinal properties of plumbagin and its analogs.” Med Res Rev 32(6): 1131–1158.

28. Poonepalli, A., L. Balakrishnan, A. K. Khaw, G. K. Low, M. Jayapal, R. N. Bhattacharjee, S. Akira, A. S. Balajee and M. P. Hande (2005). “Lack of poly(ADP-ribose) polymerase-1 gene product enhances cellular sensitivity to arsenite.” Cancer Res 65(23): 10977–10983.

29. Roake, C. M. and S. E. Artandi (2020). “Regulation of human telomerase in homeostasis and disease.” Nat Rev Mol Cell Biol 21(7): 384–397.

30. Roos, W. P. and B. Kaina (2006). “DNA damage-induced cell death by apoptosis.” Trends Mol Med 12(9): 440–450.

31. Sameni, S. and M. P. Hande (2016). “Plumbagin triggers DNA damage response, telomere dysfunction and genome instability of human breast cancer cells.” Biomed Pharmacother 82: 256–268.

32. Seimiya, H., T. Oh-hara, T. Suzuki, I. Naasani, T. Shimazaki, K. Tsuchiya and T. Tsuruo (2002). “Telomere shortening and growth inhibition of human cancer cells by novel synthetic telomerase inhibitors MST-312, MST-295, and MST-1991.” Mol Cancer Ther 1(9): 657–665.

33. Serrano, D., A. M. Bleau, I. Fernandez-Garcia, T. Fernandez-Marcelo, P. Iniesta, C. Ortiz-de-Solorzano and A. Calvo (2011). “Inhibition of telomerase activity preferentially targets aldehyde dehydrogenase-positive cancer stem-like cells in lung cancer.” Mol Cancer 10: 96.

34. Shay, J. W. and W. E. Wright (2002). “Telomerase: a target for cancer therapeutics.” Cancer Cell 2(4): 257–265.

35. Shay, J. W. and W. E. Wright (2005). “Mechanism-based combination telomerase inhibition therapy.” Cancer Cell 7(1): 1–2.

36. Smogorzewska, A., B. van Steensel, A. Bianchi, S. Oelmann, M. R. Schaefer, G. Schnapp and T. de Lange (2000). “Control of human telomere length by TRF1 and TRF2.” Mol Cell Biol 20(5): 1659–1668.

37. Sugarman, E. T., G. Zhang and J. W. Shay (2019). “In perspective: An update on telomere targeting in cancer.” Mol Carcinog 58(9): 1581–1588.

38. Tamakawa, R. A., H. B. Fleisig and J. M. Wong (2010). “Telomerase inhibition potentiates the effects of genotoxic agents in breast and colorectal cancer cells in a cell cycle-specific manner.” Cancer Res 70(21): 8684–8694.

39. van Steensel, B. and T. de Lange (1997). “Control of telomere length by the human telomeric protein TRF1.” Nature 385(6618): 740–743.

40. van Steensel, B., A. Smogorzewska and T. de Lange (1998). “TRF2 protects human telomeres from end-to-end fusions.” Cell 92(3): 401–413.

41. Venkatesan, S., A. K. Khaw and M. P. Hande (2017). “Telomere Biology-Insights into an Intriguing Phenomenon.” Cells 6(2).

42. Wong, V. C., J. Ma and C. E. Hawkins (2009). “Telomerase inhibition induces acute ATM-dependent growth arrest in human astrocytomas.” Cancer Lett 274(1): 151–159.

43. Yamada, M., N. Tsuji, M. Nakamura, R. Moriai, D. Kobayashi, A. Yagihashi and N. Watanabe (2002). “Down-regulation of TRF1, TRF2 and TIN2 genes is important to maintain telomeric DNA for gastric cancers.” Anticancer Res 22(6A): 3303–3307.

44. Zeegers, D., S. Venkatesan, S. W. Koh, G. K. Low, P. Srivastava, N. Sundaram, S. Sethu, B. Banerjee, M. Jayapal, O. Belyakov, R. Baskar, A. S. Balajee and M. P. Hande (2017). “Biomarkers of Ionizing Radiation Exposure: A Multiparametric Approach.” Genome Integr 8: 6.

